# Pathology dynamics in healthy-toxic protein interaction and the multiscale analysis of neurodegenerative diseases

**DOI:** 10.1101/2021.02.27.433219

**Authors:** Swadesh Pal, Roderick Melnik

## Abstract

Neurodegenerative diseases are frequently associated with aggregation and propagation of toxic proteins. In particular, it is well known that along with amyloid-beta, the tau protein is also driving Alzheimer’s disease. Multiscale reaction-diffusion models can assist in our better understanding of the evolution of the disease. We have modified the heterodimer model in such a way that it can now capture some of critical characteristics of this evolution such as the conversion time from healthy to toxic proteins. We have analyzed the modified model theoretically and validated the theoretical findings with numerical simulations.

## 1 Introduction

Alzheimer’s disease (AD) is an example of a neurodegenerative disease, associated with aggregation and propagation of toxic proteins [1]. Initially, the “amyloid cascade hypothesis” has dominated for the treatments [2, 3]. However, due to the failures of large clinical trials, researchers started focussing on some other mechanisms. It is now well accepted that the tau-protein (*τ P*) is a viable alternative to the “amyloid cascade hypothesis”.

The *τ P* plays a prominent role as a secondary agent in the disease development. For example, (i) frontotemporal lobar degeneration is mostly dominated by *τ P* spreading [4], (ii) neurofibrillary tangles (NFT) are correlated in brain atrophy in AD [5, 6], (iii) lower *τ P* concentration prevents neuronal loss [7], (iv) *τ P* also reduces neural activity [8], etc. This helps to explain the relative lack of clinical improvements. There is an open debate in the literature on the roles of *Aβ* proteopathy and *τ P* tauopathy in AD but it is clear by now that “the amyloid — *β* − *τ* nexus is central to disease-specific diagnosis, prevention and treatment” [9]. In recent years, many researchers have focussed on *Aβ* and *τ P* interaction. Moreover, in neurodegenerative diseases, the protein-protein interactions become a key for understanding both spreading and toxicity of the proteins [10–13]. There are some crucial observations specific to AD [10, 14]: (i) the seeding of new toxic *τ P* is enhanced by the presence of *Aβ*, (ii) the toxicity of *Aβ* depends on the presence of *τ P*, and (iii) *Aβ* and *τ P* amplify each other’s toxic effects.

Mathematical models are widely used for the interpretation of biological processes, and this field is not an exception. Building on earlier advances [15–20], we analyze AD by a deterministic mathematical modelling approach and predict the dynamics of the disease based on several novel features of our model. Recall that the heterodimer model describes the interaction between the healthy and toxic proteins [21–25]. In the heterodimer model, with an increase in the healthy protein density, the toxic protein conversion increases. We have modified that linear conversion term to include nonlinear effects via Holling type-II functional response. In this case, the toxic protein’s conversion rate remains constant with an increase in the healthy protein density. This modification incorporates the reaction time (conversion from healthy protein to toxic protein). We have considered two modified coupled heterodimer systems for healthy-toxic interactions for both proteins, *Aβ* and *τ P*, along with a single balance interaction term. This study identifies two types of disease propagation modes depending on the parameters: primary tauopathy and secondary tauopathy.

This manuscript is organized as follows. In Section 2, we briefly discuss the heterodimer model and its modification. Temporal behaviour of the modified model is analyzed in Section 3, focussing on possible stationary points and linear stability. In Section 4, we provide results on the wave propagation described by a simplified model, specific to stationary states. Section 5 is devoted to a detailed analysis of the AD propagation in terms of primary and secondary tauopathies. Concluding remarks are given in Section 6.

## 2 Mathematical Model

We first consider the usual heterodimer model for the healthy and toxic variants of the proteins *Aβ* and *τ P*. Let *Ω* be a spatial domain in ℝ^3^. For **x** ∈ *Ω* and time *t* ∈ ℝ^+^, let *u* = *u*(**x**, *t*) and *ũ*= *ũ*(**x**, *t*) be healthy and toxic concentrations of the protein *Aβ*, respectively. Similarly, we denote *v* = *v*(**x**, *t*) and 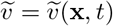 the healthy and toxic concentrations of *τ P*, respectively. The concentration evolution is then given by the following system of coupled partial differential equations [14, 22]:

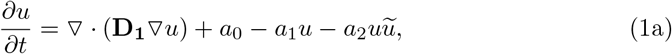

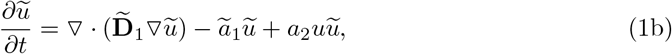

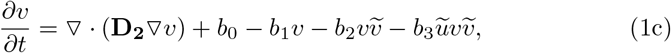

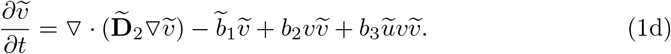

In system (1), *a*_0_ and *b*_0_ are the mean production rates of healthy proteins, *a*_1_, *b*_1_, *ã*_1_ and 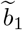 are the mean clearance rates of healthy and toxic proteins, and *a*_2_ and *b*_2_ represent the mean conversion rates of healthy proteins to toxic proteins. The parameter *b*_3_ is the coupling constant between the two proteins *Aβ* and *τ P*. Further, **D**_1_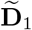, **D**_2_ and 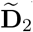 are the diffusion tensors which characterize the spreading of each proteins. We assume that all variables and initial conditions are non-negative and also all the parameters to be strictly positive.

Here, the healthy protein is approached by the toxic protein, and after transitions, a healthy protein is converted into a toxic state. In the current formulation, we have assumed that the probability of a given toxic protein encountering healthy protein in a fixed time interval *T*_*t*_, within a fixed spatial region, depends linearly on the healthy protein density. In this case, the total density of the healthy proteins *u* converted by the toxic proteins *ũ*can be expressed as *ũ*= *aT*_*s*_*u*, following the Holling functional response idea [26]. The parameter *T*_*s*_ is the time to getting contact with each other and *a* is a proportionality constant. If there is no reaction time, then *T*_*s*_ = *T*_*t*_ and hence we get a linear conversion rate *ũ*= *aT*_*t*_*u*. Now, if each toxic protein requires a reaction time *h* for healthy proteins that are converted, then the time available to getting contact becomes *T*_*s*_ = *T*_*t*_ − *hũ*. Therefore, *ũ*= *a*(*T*_*t*_ *hu*)*u*, hence *ũ*= *aT*_*t*_*u/*(1 + *ahu*), which is a nonlinear conversion rate. So, we modify the above model (1) as follows:

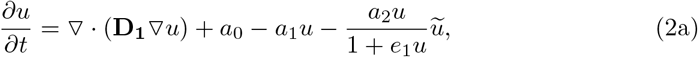

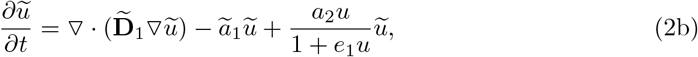

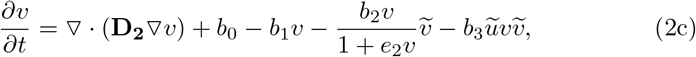

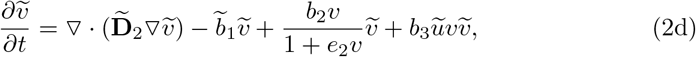

where *e*_1_ (= *a*_*β*_*h*_*β*_) and *e*_2_ (= *a*_*τ*_ *h*_*τ*_) are dimensionless parameters. We use no-flux boundary conditions and non-negative initial conditions. Here, in model (2), the rate of conversion of the healthy protein by the toxic protein increases as the healthy protein density increases, but eventually it saturates at the level where the rate of conversion remains constant regardless of increases in healthy protein density. On the other hand, in model (1), the rate of conversion of the healthy protein by the toxic protein rises constantly with an increase in the healthy protein density.

## 3 Temporal Dynamics

For studying the wave propagation based on the reaction-diffusion model (2), we will first find homogeneous steady-states of the system. The homogeneous steady-states of the system (2) can be determined by finding the equilibrium points of the following system

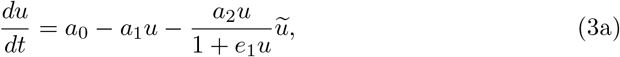

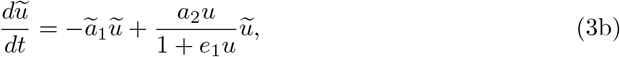

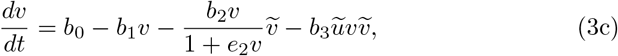

c

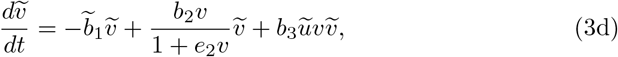

with non-negative initial conditions.

### 3.1 Stationary Points

The system (3) always has a disease-free state called a healthy stationary state. Depending on the parameter values, the system may possess more stationary points. We summarise each possible stationary state in the following:

1. Healthy *Aβ* - healthy *τ P*: It is the trivial stationary state and is given by

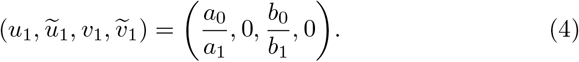

This stationary state is the same for both systems, (1) and (2), due to zero toxic loads.
2. Healthy *Aβ* - toxic *τ P*: The stationary state of “healthy *Aβ* - toxic *τ P* “ is given by

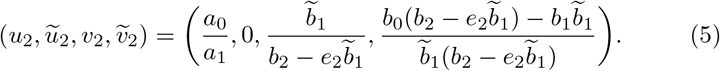

For the non-negativity of the stationary point (5), we must have 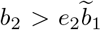 and 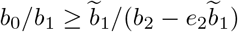.
3. Toxic *Aβ* - healthy *τ P*: The stationary state of “toxic *Aβ* - healthy *τ P* “ is given by

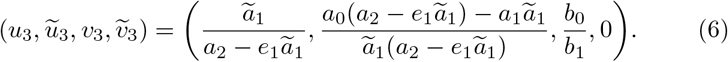

For the non-negativity of the stationary point (6), we must have *a*_2_ *> e*_1_- *ã* _1_ and *a*_0_ */a*_1_ ≥ - *ã* _1_ */*(*a*_2_ − *e*_1_ - *ã* _1_).
4. Toxic *Aβ* - toxic *τ P*: Suppose 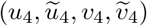 is a stationary state of the “toxic *Aβ* - toxic *τ P* “ type. In this case, we obtain *u*_4_ = *u*_3_, *ũ*_4_ = *ũ*_3_, 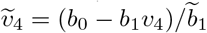 and *v*_4_ satisfy the quadratic equation

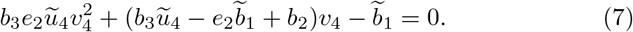

The equation (7) always has a real positive solution. For the uniqueness of *v*_4_, we must have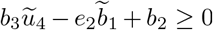. Also, for the positivity of 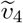, we need *v*_4_ *< b*_0_*/b*_1_.

Note that under small perturbations of any one of these stationary points, the trajectories may or may not come to that stationary point. Next, we examine this situation in more detail by the linear stability analysis.

### 3.2 Linear Stability Analysis

For the stability analysis, we linearize the system (3) about any of the stationary points 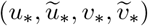. The coefficient matrix **M** of the resulting system is the Jacobian matrix of the system (3) and is given by

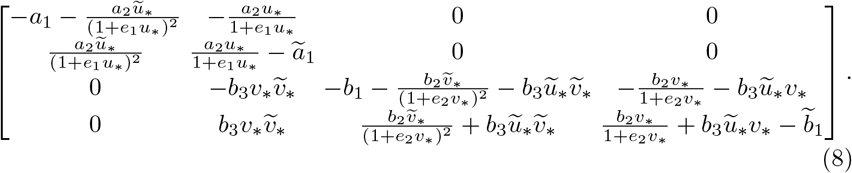

Now, the eigenvalues of the Jacobian matrix **M** are given by

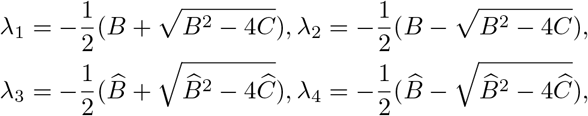

where *B* = *a*_1_ +*ã*_1_ + *a*_2_ *ũ*_∗_/(1+*e*_1_*u*_∗_)^2^ − *a*_2_*u*_∗_ /(1 + *e*_1_*u*_∗_), *C* = *a*_1_*ã*_1_ + *ã*_1_*a*_2_*ũ*_∗_/(1 + *e*_1_*u*_∗_)^2^ − *a*_1_*a*_2_*u*_*_/(1 + *e*_1_*u*_*_),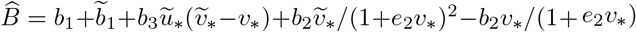 and 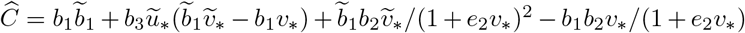.

For each of the stationary points, we find the Jacobian matrix **M** and all its eigenvalues λ_*i*_, *i* = 1, 2, 3, 4. Hence, the conclusion can be drawn easily, because for a given stationary point, if all the eigenvalues have negative real parts, this stationary point is stable, otherwise it is unstable.

## 4 Wave Propagation

We analyze travelling wave solutions of the spatio-temporal model (2) in one dimension (*Ω* = ℝ) connecting any two stationary states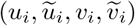, *i* = 1, 2, 3, 4 [21]. First, we consider the travelling wave emanating from healthy stationary state 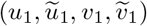 and connecting to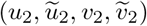. For analysing the travelling wave fronts, we linearize the spatio-temporal model (2) around the healthy stationary state which leads to the following uncoupled system

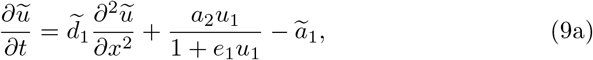

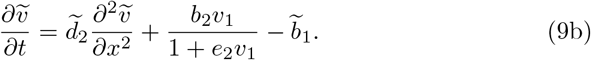

Firstly, for the travelling wave solution, we substitute *ũ*(*x, t*) = *u*(*x* − *ct*) ≡ *ũ*(*z*), 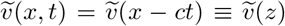 in (9) and will look for linear solutions of the form *ũ*= *C*_1_exp(*λz*),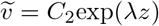. Then, the minimum wave speeds *c*_min_ are given by

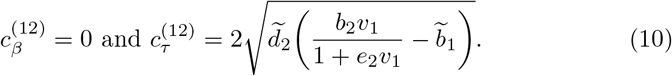

Here, 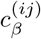 and 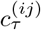 denote the speeds of the front from state *i* to the state *j* for the *Aβ* fields (*u, ũ*) and *τ P* fields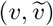, respectively.

Similarly, the minimum wave speeds for the travelling wave fronts emanating from healthy stationary state 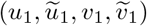 and connecting to 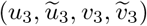 are given by

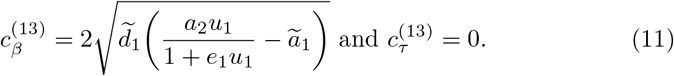

Also, we have

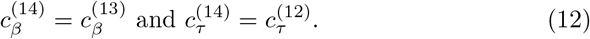

Secondly, we consider the travelling wave emanating from the stationary state 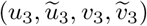 and connecting to 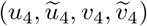. We linearize the spatio-temporal model (2) around 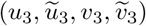 and repeat the same techniques to deduce that

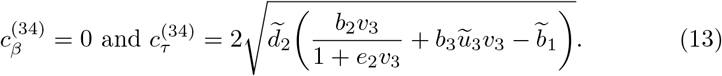

## 5 Results and Discussion

A “healthy *Aβ* - healthy *τ P* “ stationary state satisfies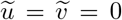. The non-existence of a physically relevant healthy state occurs due to a failure of healthy clearance, which is either with an *Aβ* clearance or with a *τ P* clearance. To start with, we consider the following balance of clearance inequalities:

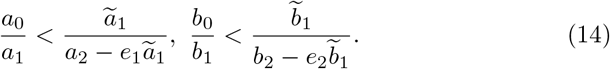

Now, if (14) holds for the stationary point 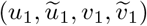 in (4), then all the eigenvalues corresponding to the Jacobian matrix **M** have negative real parts. So, given the small amounts of the production of toxic *Aβ* or toxic *τ P*, or excess amounts of the production of healthy *Aβ* or healthy *τ P*, the system would be approaching towards the “healthy *Aβ*- healthy *τ P* “ stationary state.

Due to the failure of the clearance inequality (14), a transcritical bifurcation occurs for the homogeneous system (3). Hence, all the other stationary states 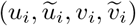, *i* = 2, 3, 4 are physically meaningful and a pathological development becomes possible. Motivated by [14], we fix the parameter values as *a*_0_ = *a*_1_ = *a*_2_ = *b*_0_ = *b*_1_ = *b*_2_ = 1 and *e*_1_ = *e*_2_ = 0.1. Now, we fix *a*_1_ = 3/4, 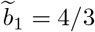 and we take *b*_3_ as the bifurcation parameter. For *b*_3_ *<* 1.575, the system has only two stationary points 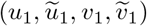 and 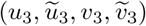. The equilibrium point 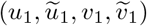is saddle and 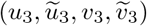 is stable. A non-trivial stationary point 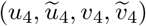 is generated through a transcritical bifurcation at *b*_3_ = 1.575. Then 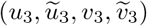 changes its stability 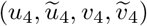 and becomes saddle (see Fig. 1).

**Fig. 1:**
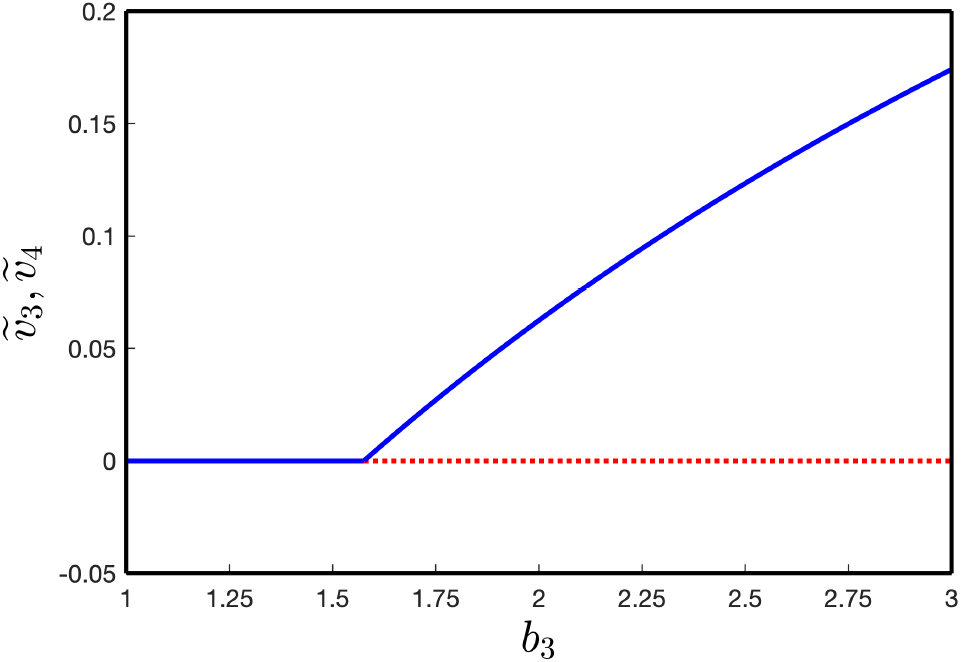
Transcritical bifurcation diagram of the stationary points for the system (3). (Parameter values: *a*_0_ = *a*_1_ = *a*_2_ = *b*_0_ = *b*_1_ = *b*_2_ = 1, *ã*_1_ = 3/4, 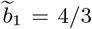, *e*_1_ = *e*_2_ = 0.1.)

These results could lead to a number of important observations. For example, due to the instability of the healthy stationary state of the system (3), a proteopathic brain patient would be progressing toward a disease state. The actual state would depend on the parameter values. If 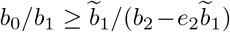 holds, then 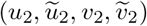 exists and if *a*_0_*/a*_1_ *ã*_1_/(*a*_2_ *e*_1_ *ã*_1_) holds, then 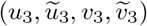 exists. Sometimes both the relations hold simultaneously. Also, the proteopathic state 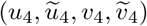 exists if *b*_0_*/b*_1_ *> v*_4_ holds. Since, *ũ*_4_ = *ũ*_3_, we can choose *b*_3_ in such a way that 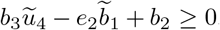. Therefore, to produce tau proteopathy, the stationary state 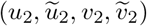 is not needed. So, we study only two types of patient proteopathies: (i) primary tauopathy and (ii) secondary tauopathy.

For the primary tauopathy, which is usually related to neurodegenerative diseases such as AD, all the four stationary states exist i.e., both the conditions 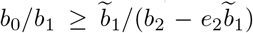 and *a*_0_*/a*_1_ *ã*_1_/(*a*_2_ *e*_1_*ã*_1_) hold. In this case, we have plotted the dynamics of the system (3) in Fig. 2(a). Also, for the secondary tauopathy, only three stationary states exist. Here, the inequality *a*_0_*/a*_1_ ≥ *a*_1_/(*a*_2_ *e*_1_*a*_1_) is true and the other inequality fails. An example of secondary tauopathy is shown in Fig. 2(b). Comparing the homogeneous systems corresponding to (1) and (2), the modified system requires less toxic load.

**Fig. 2:**
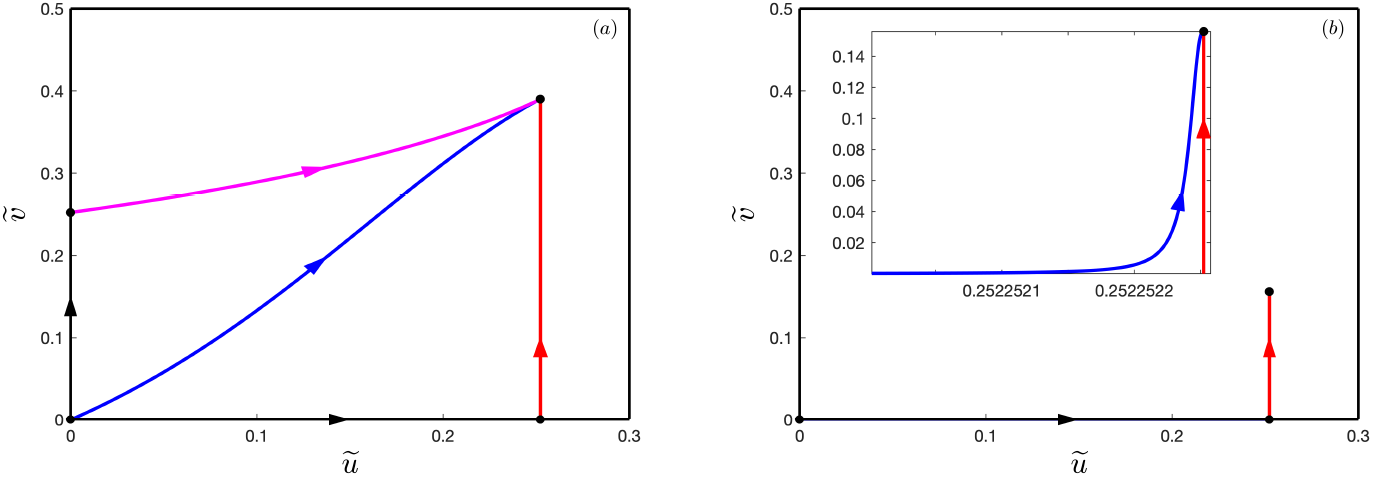
Phase plane 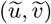 with four and three stationary points for the system (3): (a) 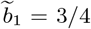, *b*_3_ = 0.5 and (b) 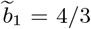, *b*_3_ = 3. (Parameter values: *a*_0_ = *a*_1_ = *a*_2_ = *b*_0_ = *b*_1_ = *b*_2_ = 1, *ã*_1_ = 3/4, *e*_1_ = *e*_2_ = 0.1.)

For the wave propagation, we consider the spatial domain in one dimension [−100, 100] as an example. However, the results are robust for a wide range of intervals. We take the initial condition 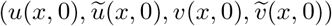 for the primary tauopathy as 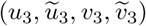 for −100 ≤ *x* ≤ −95, 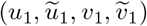 for 95 *< x <* 95 and 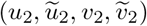 for 95 ≤ *x* ≤ 100. On the other hand, the initial condition 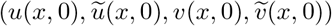 for the secondary tauopathy has been taken as 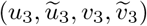 for −100 ≤ *x* ≤ −95, 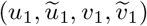 for −95 *< x <* 95 and 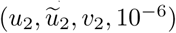 for 95 ≤ *x* ≤ 100.

We have shown the wave propagation for the primary tauopathy in Fig. 3 at different time steps *t* = 50, 150, 180 and 220. Motivated by Thompson et al., we have chosen the parameter values as *a*_0_ = *a*_1_ = *a*_2_ = *b*_0_ = *b*_1_ = *b*_2_ = 1, 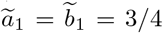, *b*_3_ = 0.5, *e*_1_ = *e*_2_ = 0.1, 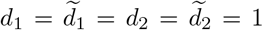 and no-flux boundary conditions for all the variables. For these parametric values, we obtain 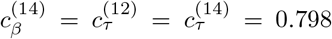 and 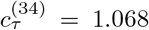. In the simulation, we have considered the toxic *Aβ* front on the left side of the domain and toxic *τ P* on the right. Initially, the toxic *Aβ* front propagates to the right with speed 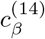 and toxic *τ P* propagates to the left with speed 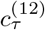. After overlapping both the fronts, *τ P* increases its concentration and connects to 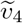. Then, the left front of the wave of *τ P* boosts its speed to 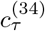 and moves to the left. On the other hand, the right front of the wave of *τ P* moves with speed 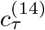, it eventually fills the domain and the entire system converges to the stable equilibrium solution 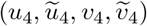.

**Fig. 3:**
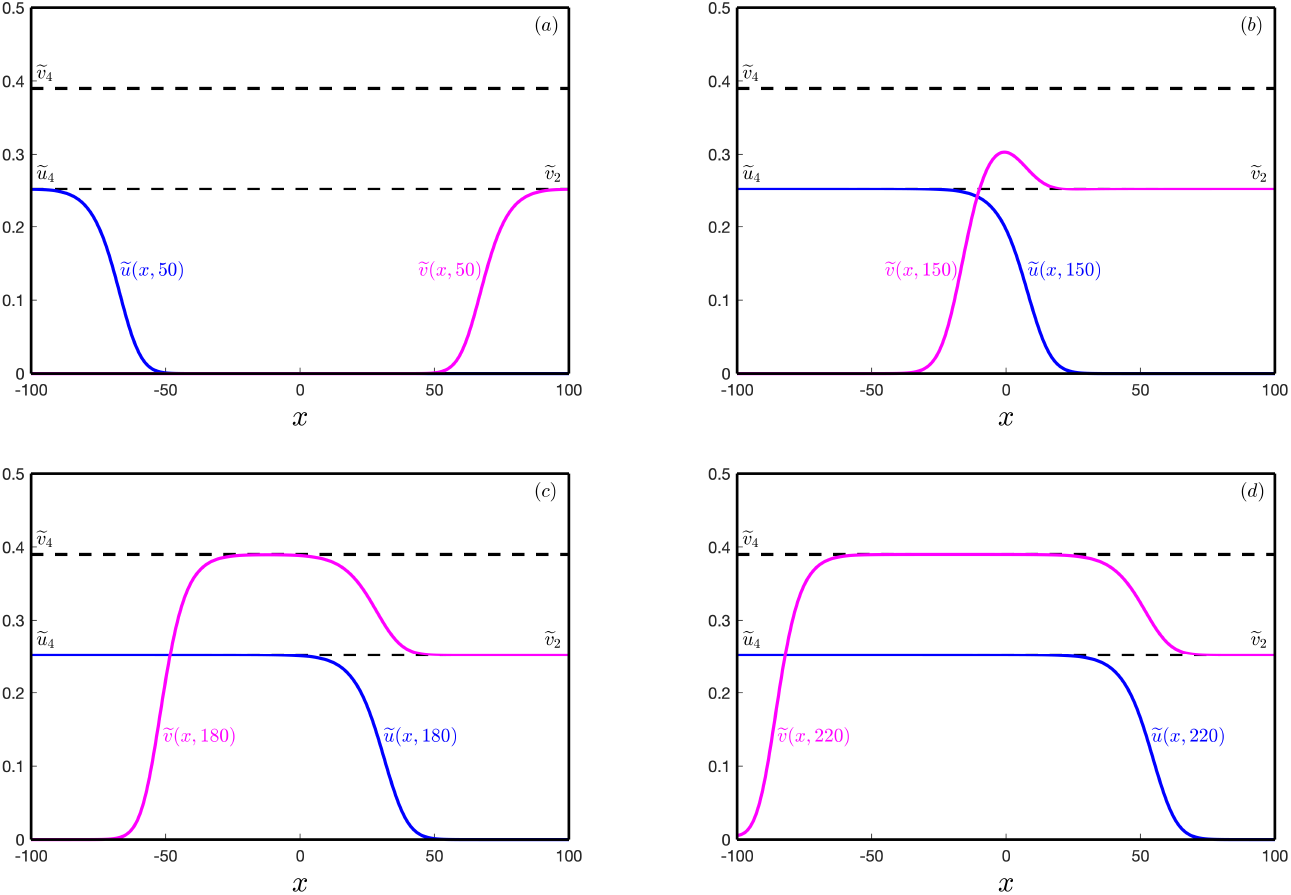
Front propagations of *ũ* and 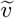 for the system (2) at different time steps: (a) *t* = 50, (b) *t* = 150, (c) *t* = 180 and (d) *t* = 220.

In Fig. 4, we plot wave propagation for the secondary tauopathy at different time steps *t* = 60, 250, 400 and 425. We have chosen the parameter values as *a*_0_ = *a*_1_ = *a*_2_ = *b*_0_ = *b*_1_ = *b*_2_ = 1, *ã*_1_ = 3/4, = 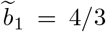, *b*_3_ = 3, *e*_1_ = *e*_2_ = 0.1, 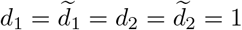 and no-flux boundary conditions for all the variables. For these parametric values, we obtain 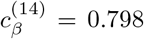 and 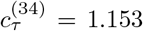. Here, the toxic *Aβ* front propagates to the right with speed 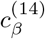 and fills the domain *ũ*_4_ with negligible toxic *τ P* (see Fig. 2(b)). However, we note that after filling toxic *Aβ* in the entire domain, toxic *τ P* starts to increase its concentration and connects to 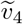. It moves with the speed 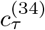 and fills the domain. Finally, the entire system converge to the stable equilibrium solution 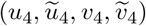.

**Fig. 4:**
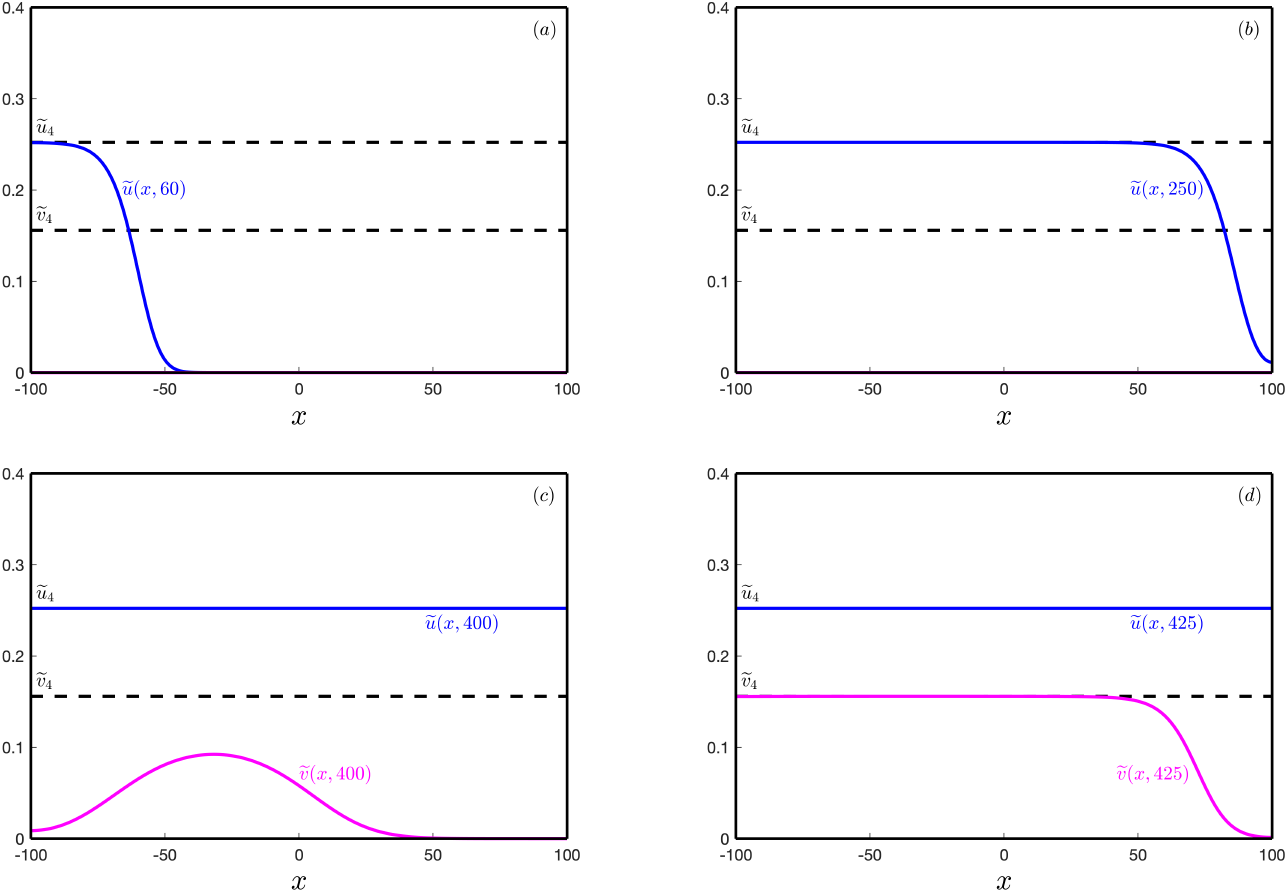
Front propagations of *ũ* and 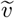 for the system (2) at different time steps: (a) *t* = 60, (b) *t* = 250, (c) *t* = 400 and (d) *t* = 425.

## 6 Conclusion

We have studied a modification of the heterodimer model, which captures the conversion time from healthy to toxic proteins. For the temporal dynamics, we have carried out the linear stability analysis of all the stationary points. We have also investigated the wave speeds of the travelling wavefronts for the spatiotemporal model.

We have obtained two clinically interesting patient proteopathies for further detailed analysis: primary and secondary tauopathies. For the case of primary tauopathy, a possible invasion of *τ P* exists independent of the invasion of *Aβ*. On the other hand, for the secondary tauopathy, the sustained presence of toxic *τ P* requires the company of toxic *Aβ*. These conclusions are similar for both the models (heterodimer and the modified version). However, for the same parametric values, the introduction of Holling type-II functional response decreases the concentrations of toxic *τ P* and toxic *Aβ* compared to the original model.

